# Correlation-resolved sum-frequency generation microscopy of biological tissue with bright twin beams

**DOI:** 10.64898/2026.10.25.761624

**Authors:** Ting-Hao Chen, Manato Adachi, Zuliang Hu, Minori Masaki, Yuriho Nikaido, Ryo Suzuki, Mutsuo Nuriya, Yujiro Eto, Hideaki Kano

## Abstract

Sum-frequency generation (SFG) driven by bright twin beams from high-gain parametric down-conversion yields a coherent component from correlated mode pairs and an incoherent component from uncorrelated mode combinations. Modal correlations can enhance coherent SFG in anisotropic nonlinear media, for example by extending the effective interaction length. Whether these correlations can also benefit image formation in biological tissue, where molecular organization and refractive index vary spatially, remains unknown. Here we address this question by performing SFG imaging of biological tissue using bright twin beams as the excitation source. Spectral decomposition yields co-registered coherent and incoherent images from a single acquisition. In mouse skeletal muscle, the coherent-component image revealed sarcomere periodicity with a Bragg-spot-to-background ratio approximately 40 times that of the incoherent-component image. In fresh human dermis, the coherent-component image showed greater Fourier power in the 0.30-0.50 μm^-1^ band and greater fractional autocovariance decay at short spatial lags than the incoherent-component image. These results demonstrate that twin-beam correlations can enhance contrast and detectability at selected spatial frequencies in heterogeneous tissue. Correlation-resolved SFG microscopy thus offers a route to label-free characterization of tissue microstructure with bright quantum light, with potential applications in biomedical research.

---

Nonlinear optical microscopy converts intrinsic molecular organization into label-free contrast. Second-order processes are particularly sensitive to non-centrosymmetric molecular assemblies, enabling second harmonic generation (SHG) and sum-frequency generation (SFG) microscopy to visualize collagen, myosin assemblies, and chiral biological structures with sub-micrometer-scale optical sectioning.^1-6^ In conventional implementations, image formation is governed by the local nonlinear susceptibility together with the intensity, spectrum, phase, polarization and focusing geometry of the illumination. Spatial information is therefore usually discussed through the optical transfer of the microscope and the nonlinear response of the specimen.

Quantum-correlated optical fields, particularly those exhibiting correlations in frequency and transverse wavevector, provide additional degrees of freedom for image formation beyond conventional illumination parameters. Quantum imaging, however, has largely developed in low-flux regimes based on single-photon detection and coincidence measurements.^7-9^ By contrast, nonlinear microscopy generally requires substantially higher photon flux. High-gain parametric down-conversion (PDC) bridges these regimes by generating bright signal–idler twin beams, also referred to as bright squeezed vacuum (BSV).^10^ These fields combine macroscopic photon flux sufficient to drive nonlinear optical processes with strong intermodal correlations that can participate directly in image formation, making them promising sources for nonlinear quantum imaging.

For nonlinear quantum imaging, a distinctive feature of twin-beam-driven SFG is the simultaneous generation of a coherent component from correlated signal–idler mode pairs and an incoherent component from interactions between uncorrelated modes. The narrow coherent spectral feature exploited here was reported as early as 1986 in frequency-doubled degenerate parametric light and was attributed to the second-order coherence and twin-photon correlation of the source, demonstrating that it can be observed without single-photon counting.^11^ Through correlations in frequency and transverse wavevector, the coherent component reconstructs the spectral and transverse-wavevector distributions of the PDC pump.^8,10,12-17^ Studies using bulk birefringent crystals have further shown that this component can mitigate group-velocity mismatch and spatial walk-off, thereby extending the effective interaction length.^10,15-17^ A quantum-optical description of these correlations is provided in Supplementary Notes 1.

This raises the question of whether the same correlation-enabled functionality can operate in biological tissue. Unlike homogeneous nonlinear crystals, biological tissue exhibits spatial variations in second-order susceptibility, molecular orientation and refractive index, which in turn affect phase accumulation, scattering and collection efficiency. It is therefore not evident that twin-beam correlations can remain operational and support a distinct coherent image channel in such a heterogeneous nonlinear medium.

Here, we establish correlation-resolved SFG microscopy by integrating our bright twin-beam source with an established multimodal nonlinear optical microscope.^18-22^ Spectral decomposition separates the coherent and incoherent SFG components, yielding two coregistered image channels from each acquisition. Measurements in skeletal muscle and dermis show enhanced contrast and detectability at selected spatial frequencies in the coherent channel, demonstrating correlation-resolved nonlinear image formation in heterogeneous tissue.

## Results

### Bright twin beams enable correlation-resolved SFG microscopy

The system combines a bright twin-beam source with sample-scanning spectral imaging, recording an SFG spectrum at each sample position (Fig. 1). A Q-switched 532-nm laser (~4 ns pulse duration, 7 kHz repetition rate) drives unseeded high-gain PDC in a temperature-controlled periodically poled lithium niobate (PPLN) crystal. The resulting signal and idler beams, centered on opposite sides of the degeneracy wavelength, each had a spectral full width at half maximum of approximately 20 nm.^23^ Residual pump light is removed using a long-pass filter. The signal and idler beams were then introduced into a custom-built sample-scanning spectroscopic microscope based on our previously reported multimodal nonlinear optical platforms.^21,22^ A 60× water-immersion objective (NA 1.27) focused the twin beams onto the specimen. Forward-propagating SFG was collected with a 40× dry objective (NA 0.6), dispersed by an imaging spectrograph and detected with a charge-coupled device (CCD) camera.

**Fig 1.**
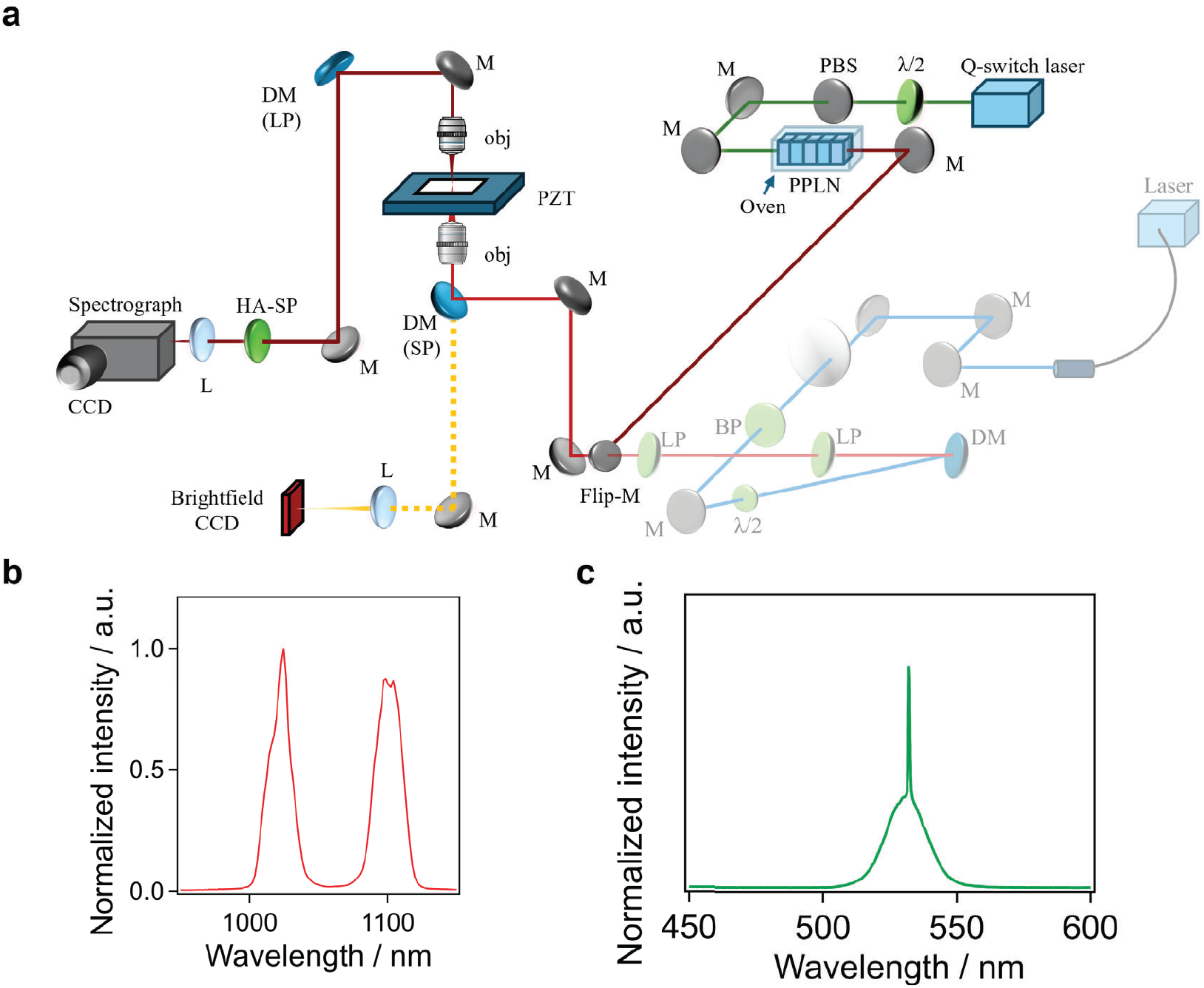
Correlation-resolved SFG microscope driven by bright twin beams. **a**, Optical layout. A 532-nm Q-switched laser pumps a temperature-controlled PPLN crystal; the generated signal and idler beams are coupled into a point-scanning microscope, and forward SFG is spectrally detected. BP, band-pass filter; DM, dichroic mirror; L, lens; LP, long-pass filter; M, mirror; obj, objective lens; PBS, polarizing beam splitter; PZT, 3-axis piezo stage; HA-SP, heat-absorbing short-pass filter; λ/2, half-wave plate. The grey optical path indicates the conventional 1064-nm excitation used for the control measurement. **b**, Representative broadband signal and idler spectrum. **c**, Representative SFG spectrum under broadband twin-beam excitation, showing a narrow pump-locked coherent feature superimposed on a broad incoherent pedestal. The sample consisted of α-quartz powder.

We decompose each SFG spectrum into a coherent component locked to the PDC pump wavelength and a broad incoherent pedestal. Let Ω_p_ be the pump angular frequency and express the signal and idler angular frequencies as ω_s_ = Ω_p_/2 + Δω and ω_i_ = Ω_p_/2 – Δω’, respectively. Correlated pairs have Δω’ = Δω and satisfy ω_s_ + ω_i_ = Ω_p_, producing the narrow, pump-locked component. Uncorrelated modes generally have Δω’ ≠ Δω, so ω_s_ + ω_i_ = Ω_p_ + Δω – Δω’ spreads over a broad range, producing the incoherent pedestal.

In a separate source-characterization experiment, both components scaled quadratically with the transmitted twin-beam intensity when the signal and idler beams were attenuated together after PDC, confirming their second-order optical origin (Supplementary Note 2 and Supplementary Fig. S1). The coherent SFG component can contain contributions proportional to *N*^2^ and *N*, where *N* is the mean photon number per mode, and the linear contribution need not be confined to the isolated-pair regime.^18,24^ Because post-PDC attenuation gives the *N*^2^ and *N* contributions the same transmission dependence, this measurement confirms the second-order optical origin but does not determine their relative weights.

### Coherent SFG resolves the sarcomere periodicity of mouse skeletal muscle

We first examined mouse gastrocnemius (lower-leg skeletal muscle), in which myosin thick filaments generate a strong second-order response and the sarcomere lattice provides an independently known periodicity. The field-averaged spectrum showed a pump-locked coherent peak superimposed on an incoherent pedestal (Fig. 2a). Pixel-wise decomposition produced coregistered coherent and incoherent images of the same field (Fig. 2b,c).

**Fig 2.**
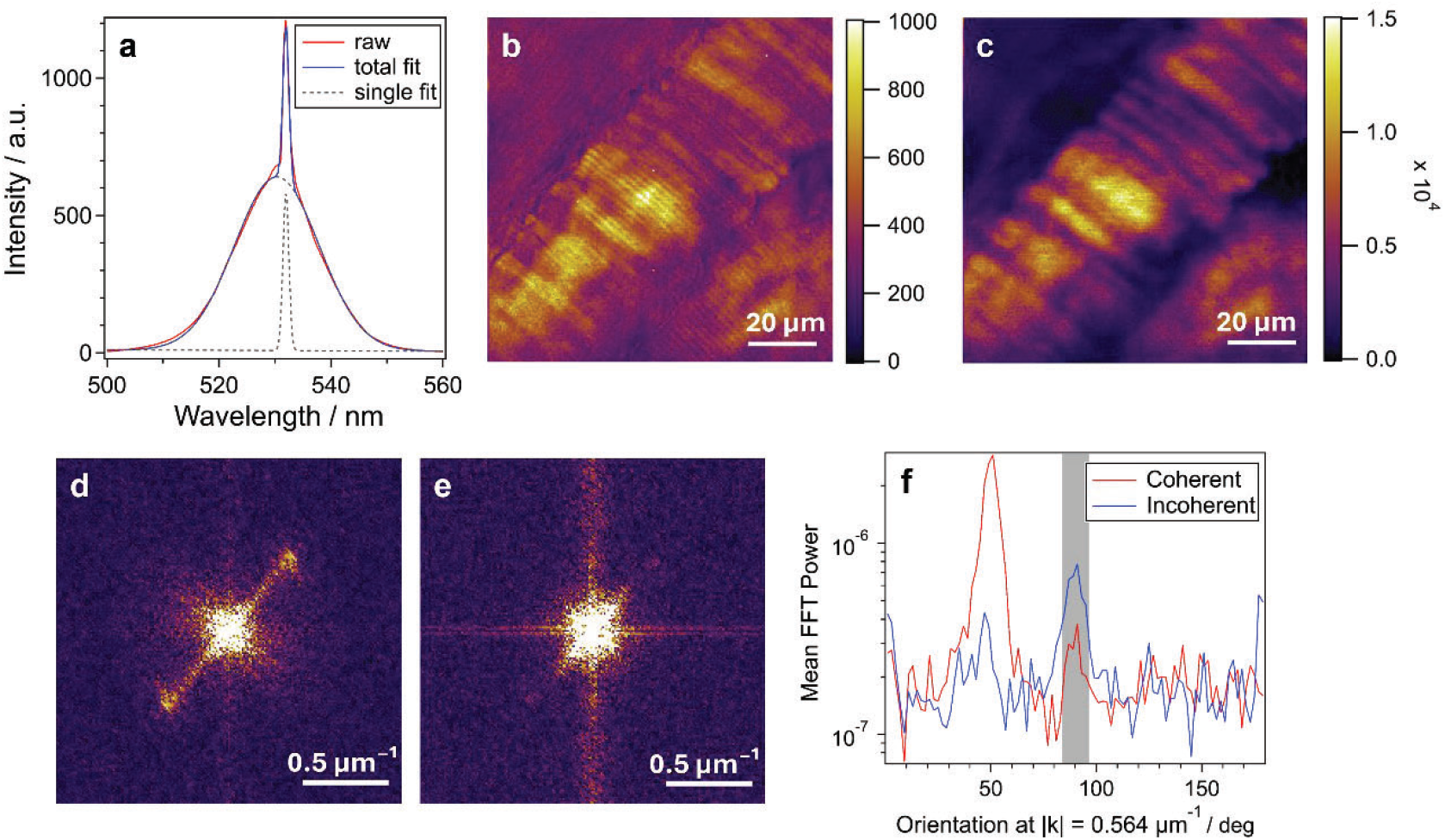
Correlation-resolved SFG resolves the sarcomere periodicity in mouse skeletal muscle. **a**, Field-averaged spectrum with the fitted coherent (sharp) and incoherent (broad) Gaussian components. The fitted constant baseline was subtracted. **b**,**c**, Images of the integrated coherent and incoherent components, respectively, reconstructed from a single hyperspectral acquisition comprising 201 × 201 pixels at a sampling pitch of 0.5 μm. Scale bars, 20 μm. **d**,**e**, Two-dimensional power spectra of b and c, displayed independently normalized to their respective maximum. The off-center features near lkl = 0.56 µm^-1^ arise from sarcomere periodicity. Scale bars, 0.5 μm^-1^. The vertical band is a fast-axis scan artifact. **f**, Fourier power at |k| = 0.56 µm^-1^ as a function of orientation, measured from the +k_γ_ direction. The grey shading marks the sectors affected by the fast-axis scan artifact.

The two images followed the same overall fiber architecture. However, sarcomere striations within individual myofibrils were clearly resolved in the coherent image but appeared only faintly in the incoherent image. Although the coherent image had a lower coefficient of variation across the field, its clearer striations reflected a larger relative contribution from high spatial frequencies rather than greater global contrast.

Fourier transformation of the images revealed a pair of Bragg spots arising from sarcomere periodicity at |k| ≈ 0.56 μm^-1^, corresponding to a period of approximately 1.8 μm and consistent with reported sarcomere lengths in mouse skeletal muscle^25^,^26^ (Fig. 2d,e). Angular profiles of the power at this spatial frequency showed discrete peaks at the Bragg-spot orientations, allowing their power to be compared with the background at the same |k| but at other orientations (Fig. 2f). The Bragg-spot-to-background ratio was approximately 40 times higher in the coherent-component image than in the incoherent-component image.

### Coherent SFG enhances contrast at selected spatial frequencies in fresh human dermis

To test whether the separation between the coherent and incoherent SFG channels persists in a structurally distinct tissue, we imaged collagen-rich fresh human dermis using the same acquisition scheme and spectral-decomposition procedure. The main field was recorded using a total signal-plus-idler power of 0.5 mW, a dwell time of 500 ms per pixel and a sampling pitch of 0.5 μm. The field-averaged spectrum again showed a narrow coherent peak superimposed on a broad incoherent pedestal (Fig. 3a). Pixel-wise spectral decomposition separated the two components across the field, yielding coherent and incoherent images (Fig. 3b,c). The dermal second-order response most likely originates from ordered collagen, whose density, orientation and local optical environment vary across the tissue. Despite this heterogeneity, both the coherent and incoherent images delineated individual collagen fiber bundles (Fig. 3b,c), with their differences emerging primarily at higher spatial frequencies.

**Fig 3.**
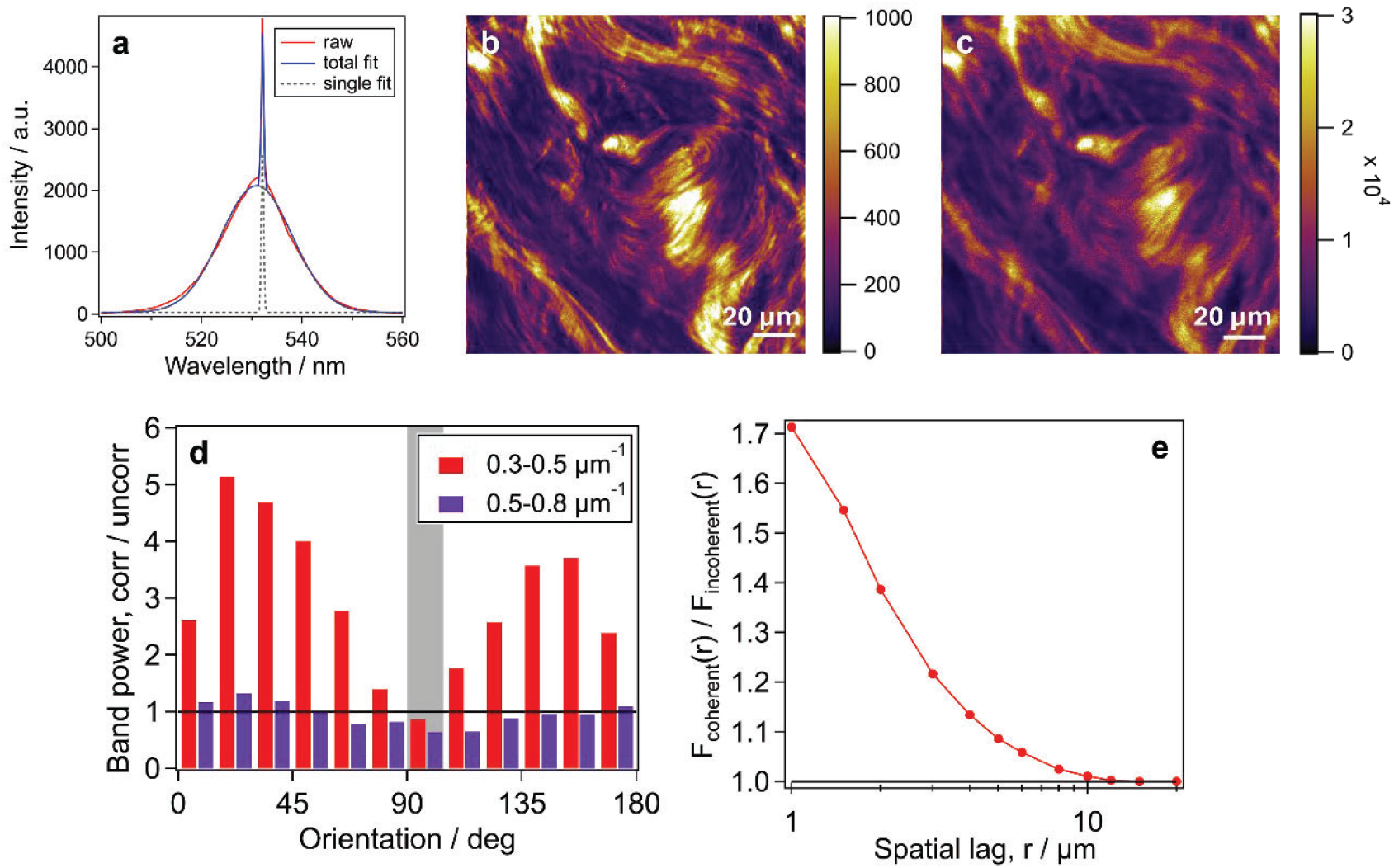
Correlation-resolved SFG imaging of fresh human dermis. **a**, Field-averaged spectrum with fitted coherent (sharp) and incoherent (broad) Gaussian components. The fitted constant baseline was subtracted. **b**,**c**, Images of the integrated coherent and incoherent components, respectively, reconstructed from a single acquisition comprising 321 × 321 pixels at a sampling pitch of 0.5 μm. Scale bars, 20 μm. **d**, Coherent-to-incoherent band-power ratio in 15° orientation sectors for the indicated spatial-frequency bands. The horizontal line marks unity. The grey shading marks the sectors affected by the fast-axis scan artifact. **e**, Coherent-to-incoherent ratio of the autocovariance decay, *F*_coherent_(r)/*F*_incoherent_(r), as a function of spatial lag r. *F*(r) is the fraction of the autocovariance decrease between the one-pixel lag and 20 μm that occurs by lag r. Values above unity indicate that a greater fraction of this decrease occurs by lag r in the coherent image than in the incoherent image. The ratio equals unity at 20 μm by construction.

To test whether the broad pedestal originated from the tissue itself, we imaged the same lateral field of the dermis with a 1064-nm laser through the same microscope. Conventional excitation produced a 532-nm SHG peak with a spectral width consistent with the bandwidth of the 1064-nm laser and no broad pedestal, supporting assignment of the broad pedestal to the broadband twin-beam excitation (Supplementary Note 3 and Supplementary Fig. S2). A small unintended axial offset between the measurements and the use of different input polarizations precluded quantitative comparison of the images.

Over the range 0.30–0.50 μm^-1^, the mean-normalized coherent image carried 1.4–5.1 times the Fourier power of the incoherent image across all orientations, with a mean ratio of 3.15 (Fig. 3d). To assess how image contrast was distributed across spatial scales, we calculated the fractional autocovariance decay, *F*(*r*), which quantifies the fraction of the autocovariance decrease from the one-pixel lag to 20 μm that has occurred by lag *r* (Methods). The coherent-to-incoherent ratio of *F*(*r*) exceeded unity at short lags and reached unity at 20 μm by construction (Fig. 3e), indicating that a greater fraction of the total autocovariance decrease occurred at short lags in the coherent image. This behavior is consistent with a greater relative contribution from fine-scale intensity variations. Together, the Fourier and autocovariance analyses show that coherent SFG provides greater micrometer-scale structural contrast in heterogeneous dermal tissue.

Finally, the coherent and incoherent SFG components were also separated across a 3 × 3 mm field in mouse tail annulus fibrosus. Both component images delineated the collagen fascicle architecture, showing that the two spectral channels can be mapped over millimeter-scale tissue fields (Supplementary Fig. S3).

## Discussion

We show that a bright twin-beam source enables two spectrally separable, co-registered SFG image channels in heterogeneous biological tissue. In skeletal muscle, the coherent channel revealed the independently known sarcomere periodicity with substantially greater Bragg-spot contrast than the incoherent channel. In fresh human dermis, the coherent channel carried greater Fourier power across all orientations in the 0.30–0.50 μm^-1^ band and exhibited a greater fractional autocovariance decay at short spatial lags. The coherent and incoherent components were also recovered throughout a 3 × 3 mm field in mouse tail annulus fibrosus. The central advance is therefore not simply the detection of a narrow spectral feature, but the formation of a distinct spatial image channel through the recombination of correlated signal–idler pairs within the tissue itself.

One possible mechanism underlying the difference in spatial contrast between the coherent and incoherent images is correlation-selective longitudinal buildup of the SFG field. Previous studies of PDC followed by SFG in bulk nonlinear crystals have shown that correlated signal–idler pairs can satisfy the corresponding SFG phase-matching condition and add coherently along the interaction length, whereas combinations of uncorrelated modes generally accumulate different longitudinal phases and dephase more rapidly.^15^,^17^ In the idealized limit, the coherent intensity increases quadratically with the effective interaction length, whereas the incoherent contribution increases approximately linearly.^17^ Spatial variations in molecular orientation, birefringence, effective interaction length and phase accumulation could therefore weight the two components differently across the tissue, potentially enhancing fine-scale contrast in the coherent image. This interpretation remains to be tested directly in tissue

This proposed longitudinal mechanism differs from the direct effect of transverse-wavevector pairing. In the type-I bulk-crystal geometry analyzed by Jedrkiewicz *et al*.,^17^ conjugate modes forming the coherent component satisfy **q**_i_ = -**q**_s_, so the transverse phase factor exp[i(**q**_s_ + **q**_i_)·**ρ**] is unity at the field-amplitude level, irrespective of |**q**_s_|. Here, **ρ** denotes the transverse coordinate in the focal plane. For the incoherent component, nonconjugate mode combinations carry different transverse phases, but ensemble averaging eliminates the cross terms between uncorrelated combinations. Only the individual intensities remain, for which |exp[i(**q** _s_ + **q**_i_)·**ρ**]|^2^ = 1. Thus, the transverse phase factor is eliminated through different mechanisms in the two components and does not introduce spatial-frequency-selective weighting into either component. This difference alone therefore cannot explain the greater high-spatial-frequency power or the greater fractional autocovariance decay at short lags observed in the coherent image. How transverse-wavevector correlations enter the walk-off-dependent effective phase mismatch within this type-I bulk-crystal model is considered further in Supplementary Note 4.

Regardless of the underlying mechanism, neither the greater high-spatial-frequency power nor the greater fractional autocovariance decay at short lags should be interpreted as evidence of an extended spatial-frequency cutoff. Establishing whether correlation-resolved SFG alters spatial resolution will require calibrated measurement of the complex amplitude spread function and the corresponding coherent transfer function. Instead, our results show that the multimode pair structure of bright twin beams provides an additional degree of freedom for image formation. Correlated mode pairs and uncorrelated mode combinations generate spectrally distinct SFG channels with different spatial contrast at macroscopic photon flux.

A practical strength of correlation-resolved SFG is its operation at macroscopic photon flux in the heterogeneous optical environment of biological tissue. The coherent and incoherent components can be separated spectrally at each pixel without single-photon detection or coincidence measurements. Their recovery in three tissues with distinct architectures, over spatial scales ranging from micrometer-scale sarcomere structure to a 3-mm tissue field, supports the robustness of the approach. Future measurements that systematically vary tissue thickness, molecular orientation, focal position, signal–idler delay and collection geometry will be needed to determine the origin and generality of the spatial differences. More broadly, correlation-resolved SFG uses the correlation structure of the illumination as an additional degree of freedom for nonlinear optical image contrast. This approach offers a basis for label-free studies of tissue organization and remodeling, with potential applications to the characterization of disease-associated changes in collagen and muscle architecture.

## Methods

### Bright twin-beam source

The bright twin beams were generated by unseeded high-gain PDC in a temperature-controlled PPLN crystal pumped by a Q-switched 532-nm laser.^23^ The pump pulse duration was 3.9 ns and the repetition rate was 7 kHz. For the tissue-imaging experiments, the signal and idler spectra centered at approximately 1022 and 1101 nm, respectively. Residual 532-nm pump light was removed after the crystal, and the twin beams were directed to the microscope. The source architecture was based on our previously reported bright twin-beam systems.^18–20^

### Correlation-resolved SFG microscope

The bright twin-beams were introduced into a custom-built multimodal nonlinear optical microscope adapted from our previously reported platforms for coherent anti-Stokes Raman scattering, SHG, third-harmonic generation and third-order sum-frequency generation imaging.^20^ A 60× water-immersion objective (NA 1.27; PlanApo, Nikon) focused the twin beams onto the specimen. Forward-propagating SFG signal was collected using a 40× dry objective (NA 0.6; SFluor, Nikon), spectrally dispersed with a spectrograph (SpectraPro 300i, Princeton Instruments) and detected with a CCD camera (PIXIS 100B, Princeton Instruments). The specimen was raster-scanned using a three-axis piezoelectric stage (LP300, Mad City Labs), with one spectrum recorded at each pixel.

The PPLN output plane was not relayed onto the specimen using a dedicated 4*f* imaging system. Instead, the twin beams were coupled into the microscope and focused directly onto the specimen by the illumination objective. For all main-text measurements, the total signal-plus-idler power incident on the specimen was approximately 0.5 mW. The exposure time was 700 ms per pixel for mouse skeletal muscle, 500 ms per pixel for fresh human dermis and 100 ms per pixel for mouse tail anulus fibrosus. The independent dermal field shown in Supplementary Fig. S2 was recorded at 0.4 mW with an exposure of 200 ms per pixel.

### Tissue preparation

C57BL/6N wild-type male mice were housed under controlled temperature and humidity on a 12-hour light/12-hour dark cycle. At 10–20 weeks of age, mice were deeply anesthetized with isoflurane, followed by intraperitoneal administration of somnopentyl, and then transcardially perfused with phosphate-buffered saline (PBS). No fixation was performed to avoid disruption of the tissue ultrastructure. Gastrocnemius (lower-leg muscle) and tail samples were collected, cryoprotected in 30% sucrose in PBS at 4°C for 1 day, embedded in OCT compound (Sakura Finetek) the following day, and frozen at -80°C. The muscle and tail samples were then cryosectioned at thicknesses of 40 and 100 µm, respectively. Microscopic observation was performed immediately thereafter. All animal experimental procedures were approved (approved protocol ID #24-324) and conducted following the guidelines established by the Institutional Animal Care and Use Committee of University of Tsukuba.

Fresh, unfixed human skin specimens were prepared and provided by Shiseido Co., Ltd. The dermis was dissected from each specimen and was estimated to be several hundred micrometers thick. Each sample was mounted between a glass slide and a coverslip and imaged without fixation. The tissues were collected during plastic surgery after informed consent had been obtained, in accordance with French law and applicable ethical regulations. Their use in this study was approved by the Ethics Committee of Shiseido and the Faculty and Graduate School of Science and Technology Research Ethics Committee of Keio University (approval no. 2026-055), in accordance with relevant institutional and national guidelines, including those of the National Institutes of Health.

### Image reconstruction and spatial-frequency analysis

For each specimen, the coherent- and incoherent-component images were reconstructed on the same spatial grid from a single hyperspectral acquisition, eliminating inter-scan registration errors. Mouse skeletal muscle was scanned over 100 × 100 μm at a pitch of 0.5 μm, producing 201 × 201 pixels. Fresh human dermis was scanned over 160 × 160 μm at the same pitch, producing 321 × 321 pixels. Both datasets therefore had a spatial Nyquist frequency of 1.0 μm^-1^. The mouse tail anulus fibrosus was imaged over a 3 × 3 mm field using an LP-200 piezoelectric stage (Mad City Labs) and our previously described seamless tiling procedure, without tile-wise intensity correction.^22^ The resulting dataset was processed using the same spectral-decomposition procedure but was excluded from the quantitative spatial-frequency comparison because of its coarser spatial sampling. That field was sampled at a 5 μm pitch, producing 600 × 600 pixels and a spatial Nyquist frequency of 0.1 μm^-1^, a tenth of that of the muscle and dermal scans; this is the coarser sampling referred to above. The pixel dwell time was 100 ms.

### Spectral decomposition

Each pixel spectrum was fitted with a constant baseline and two Gaussian components. The centers of both Gaussians and the width of the sharp Gaussian were fixed to values obtained from the field-averaged spectrum. The amplitudes of both Gaussians, the baseline, and the width of the broad Gaussian were fitted independently at each pixel. Component images were defined from the integrated areas of the fitted Gaussians. Because the sharp-component width was fixed, its area was proportional to its fitted amplitude, whereas the fitted width was included in calculating the broad-component area.

### Fourier analysis

Before Fourier transformation, each image was divided by its mean and mean-subtracted, allowing the spatial contrast of the two channels to be compared independently of their absolute fitted amplitudes. A separable Hann window was then applied, followed by a two-dimensional discrete Fourier transform without zero padding.

The spatial-frequency interval was 1/(*Nx*), where *N* is the number of pixels along one dimension and *x* is the pixel pitch. The intervals were 0.010 µm-1 for muscle and 0.0062 µm-1 for dermis, with a Nyquist frequency of 1.0 µm-1 for both datasets. Because an artifact along the fast scan axis affected the two channels differently, orientation-resolved quantities were used as the primary measures.

Discrete Fourier features were compared with the background at the same spatial-frequency magnitude, |k|, at orientations separated by more than 15° from the feature. Orientation-resolved band power was calculated in 15° sectors, both across all orientations and after excluding orientations within ±25° of the fast scan axis. The root-mean-square modulation was calculated in overlapping 20 × 20 μm (40 × 40 pixel) windows at all positions for which the full window lay within the image, without padding, after band-pass filtering around the sarcomere wavevector. Four control directions at the same spatial-frequency magnitude |k| were processed identically. Sarcomere modulation was considered detectable when the local modulation amplitude at the Bragg-spot pair exceeded three times the mean amplitude of the four controls.

### Autocovariance analysis across spatial scales

To quantify how image contrast was distributed across spatial scales, we analyzed the radial autocovariance functions of the coherent and incoherent SFG images. The autocovariance was calculated as a function of spatial lag and radially averaged to obtain *C*(*r*), where *r* denotes the spatial separation between pixels. For each component, the fractional autocovariance decay at lag *r*, where *r*_1_< *r* ≤ S *r*_ref_, was defined as

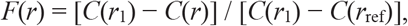

where *r*_1_ = 0.5 µm is the smallest nonzero spatial lag, corresponding to the separation between adjacent pixels, and *r*_ref_ = 20 µm is the reference lag. Thus, *F*(*r*) quantifies the fraction of the autocovariance decrease between the one-pixel lag and 20 μm that occurs by lag *r*. The fractional autocovariance decay was calculated independently for the coherent and incoherent components, and their ratio was defined as

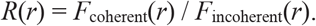

Values of *R*(*r*) > 1 indicate that a greater fraction of the autocovariance decrease over the reference range occurs by lag *r* in the coherent component than in the incoherent component. All calculations were performed using custom procedures implemented in Igor Pro (WaveMetrics).

## Statistics and reproducibility

The quantitative analyses reported here are descriptive comparisons between simultaneously reconstructed image channels within individual fields of view. A separately acquired dermal field was analyzed as an acquisition-level consistency check (Supplementary Fig. S3). No statistical inference across independent biological replicates was performed.

## Data availability

The data supporting the findings of this study are available from the corresponding author upon reasonable request.

## Code availability

The code used for spectral decomposition, image reconstruction and spatial-frequency analysis is available from the corresponding author upon reasonable request.

## Acknowledgements

The authors thank Dr. M. Egawa, Mr. K. Koizumi and Dr. C. Knight (Shiseido Co., Ltd.) for their support and valuable contributions.

## Author contributions

Y.E. and H.K. conceived the study. T.-H.C. and M.A. developed the microscope and performed the experiments. T.-H.C. and M.A. analyzed the data with contributions from Z.H., M.N., Y.N. and R.S. M.M. contributed biological expertise and resources. Y.E. developed the twin-beam source and provided guidance on correlation measurements. Y.E. and H.K. jointly supervised the project. T.-H.C., Y.E. and H.K. wrote the manuscript with input from all authors. All authors reviewed the manuscript.

