## Supplemental Document and Figures for "Correlation-resolved sum-frequency generation microscopy of biological tissue with bright twin beams"

**Supplementary Note 1 | Joint frequency–transverse-wavevector correlations of the bright PDC source**

In a sum-frequency generation (SFG) process driven by twin beams, the frequency correlation between the signal and idler modes enhances the generation of the SFG component whose spectrum corresponds to that of the PDC pump. Assuming a monochromatic PDC pump with angular frequency $\Omega_p$, the angular frequencies $\omega_s$and $\omega_i$of a correlated signal–idler mode pair satisfy

$$\omega_s + \omega_i = \Omega_p.$$

This component is commonly referred to as the coherent SFG component. An analogous effect occurs for the transverse wavevectors. Namely, if the transverse wavevectors of the signal and idler modes are denoted by $\boldsymbol{q}_s$ and $\boldsymbol{q}_i$, respectively, the generation efficiency of the SFG component satisfying

$$\boldsymbol{q}_s + \boldsymbol{q}_i = 0$$

is enhanced. The simultaneous manifestation of frequency and transverse-wavevector correlations in the coherent SFG response has been theoretically described in Ref. [1]. In the following, we describe its physical origin.

The annihilation operator of the PDC output field can be expressed using the Bogoliubov transformation as

$$\hat{a}(\boldsymbol{q},\omega) = \mu(\boldsymbol{q},\omega)\hat{a}_{\mathrm{vac}}(\boldsymbol{q},\omega) + i\nu(\boldsymbol{q},\omega)\hat{a}^{\dagger}_{\mathrm{vac}}\big(-,\boldsymbol{q}\Omega_p - \omega\big).$$

Here, $\mu(\boldsymbol{q},\omega)$ and $\nu(\boldsymbol{q},\omega)$ are the Bogoliubov coefficients. The input vacuum-field operators $\hat{a}_{\mathrm{vac}}$ and $\hat{a}^{\dagger}_{\mathrm{vac}}$ satisfy the commutation relation.

$$\big[\hat{a}_{\mathrm{vac}},(\boldsymbol{q},\omega),\ \hat{a}^{\dagger}_{\mathrm{vac}}(\boldsymbol{q},\omega)\big] = \delta^{(2)}(\boldsymbol{q}-\boldsymbol{q}')\delta(\omega-\omega').$$

The field operator of the generated SFG in the frequency domain is expressed as

$$\hat{a}_{\mathrm{SFG}}(\boldsymbol{q}_{\mathrm{SFG}},\ \omega_{\mathrm{SFG}}) \propto \int d^2\, q_s d^2 q_i \int d\,\omega_s d\omega_i\ \hat{a}(\boldsymbol{q}_s,\omega_s)\hat{a}(\boldsymbol{q}_i,\omega_i)$$

$$\times\,\delta^{(2)}(q_s + q_i - \boldsymbol{q}_{\mathrm{SFG}})\delta(\omega_s + \omega_i - \omega_{\mathrm{SFG}}).$$

The SFG intensity is given by $I_{\mathrm{SFG}} \propto \langle 0|\hat{a}^{\dagger}_{\mathrm{SFG}}|\hat{a}_{\mathrm{SFG}}0\rangle$, where $|0\rangle$ denotes the vacuum state. The integrand contains the following term:

$$\big\langle \hat{a}^{\dagger}(\tilde{\boldsymbol{q}}_i,\tilde{\omega}_i)\hat{a}^{\dagger}(\tilde{\boldsymbol{q}}_s,\tilde{\omega}_s)\hat{a}(\boldsymbol{q}_s,\omega_s)\hat{a}(\boldsymbol{q}_i,\omega_i)\big\rangle,$$

where the variables with tildes denote those associated with the ket vector.

Evaluating this expectation value using the commutation relations of the vacuum fields, we obtain a term containing the following delta functions,

$$\delta^{(2)}(\boldsymbol{q}_s + \boldsymbol{q}_i)\ \delta\big(\omega_s + \omega_i - \Omega_p\big),$$

which gives rise to the coherent component. The product of delta functions indicates simultaneous correlations in frequency and transverse wavevector.

**Supplementary Note 2 | Intensity dependence of the two spectral components**

We measured the intensity dependence of both spectral components using a 0.5-mm-thick β-barium borate (BBO) crystal as the SFG medium to confirm their nonlinear origin. With the spectrometer configured for the near infrared, the signal and idler were centered near 1024 and 1102 nm. The 532-nm upconverted SFG signal appeared in second diffraction order at 1064 nm, allowing the signal, idler and SFG to be recorded in the same frame (Supplementary Fig. S1a).

A motorized neutral-density disc provided 301 attenuation steps spanning a factor of 6.4 in signal and idler intensities. At each step, the SFG spectrum was fitted with a constant baseline and two Gaussians whose centers and widths were fixed from the summed spectrum (full widths at half maximum, 0.24 and 8.1 nm for the sharp and broad components, respectively).

Power-law fits to the integrated signal intensity gave exponents of 1.95 ± 0.02 for the coherent component and 1.84 ± 0.02 for the incoherent component (Supplementary Fig. S1b). Both components therefore showed the same nonlinear scaling within uncertainty, supporting their assignment as SFG signals.

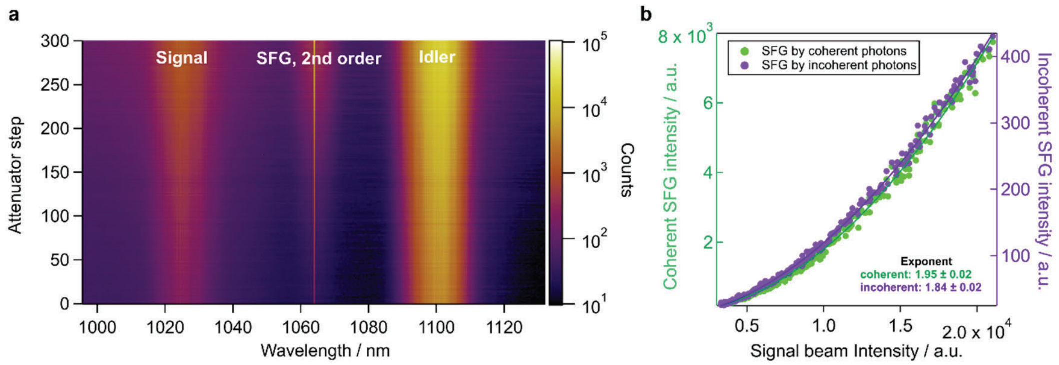


**Supplementary Fig. S1 | Intensity dependence of the coherent and incoherent spectral components.**

**a,** Attenuation-series spectra on a logarithmic color scale. Signal and idler flank the degeneracy point, and second-order diffraction places the 532-nm SFG signal at 1064 nm. **b,** Integrated coherent and incoherent SFG components plotted against the integrated signal intensity, with power-law fits. The fitted exponents were 1.95 ± 0.02 and 1.84 ± 0.02 for the coherent and incoherent components, respectively.

**Supplementary Note 3 | Comparison with a conventional laser at matched power and dwell time**

To compare excitation by bright twin beams with conventional narrowband excitation, we imaged the same lateral field of a dermal specimen through the same microscope and spectrograph using approximately 0.4 mW of incident power and a dwell time of 200 ms per pixel. Conventional 1064-nm excitation produced only a narrow 532-nm SHG peak, whereas excitation with bright twin beams produced a coherent pump-locked peak superimposed on a broad incoherent pedestal, supporting assignment of the pedestal is specific to the broadband twin-beam excitation. Signal strengths were quantified as fitted Gaussian areas. Relative to the laser peak, the coherent BSV component was 3.2-fold greater, and the total BSV signal, including both coherent and incoherent components, was 84-fold greater (Supplementary Fig. S2). Because a small axial offset remained and the input polarizations differed, these ratios are acquisition-specific rather than calibrated enhancement factors.

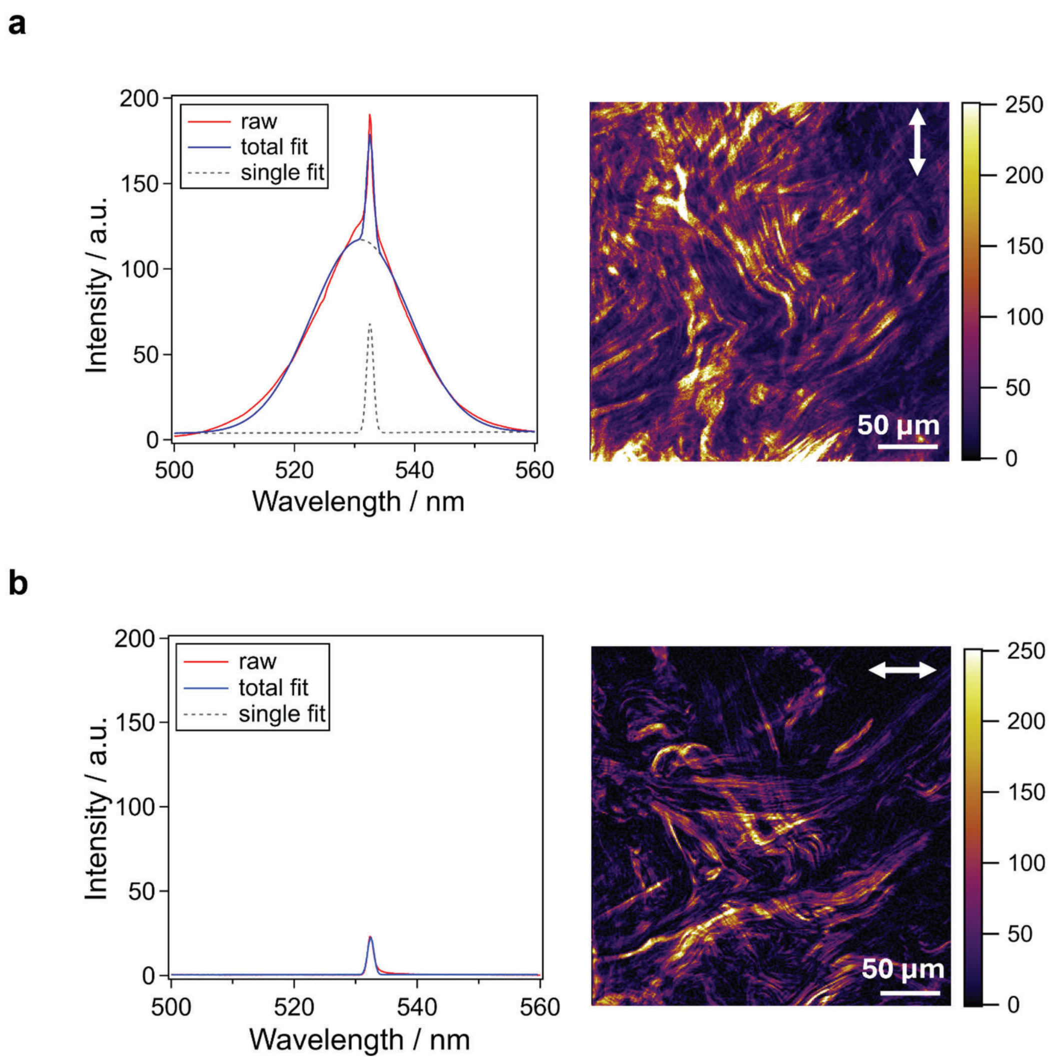


**Supplementary Fig. S2 | Comparison of field-averaged SFG/SHG spectra at matched incident power and pixel dwell time.**

a,b, Spectra acquired from the same dermal specimen using the bright twin-beam source (a) and a conventional 1064-nm laser (b). The incident power was approximately 0.4 mW and the dwell time was 200 ms per pixel. Red, measured spectrum; blue, total fit; gray dashed, fitted narrow component. Because no broad pedestal was detected in b, the laser spectrum was fitted with a single Gaussian. The same vertical scale is used in both panels. Right shows representative tissue images acquired under the corresponding excitation conditions; white arrows indicate the input polarization directions. The same lateral field was imaged in both measurements, although a small unintended axial offset remained and the input polarizations differed; the insets are therefore shown for qualitative comparison only; scale bars, 50 μm.

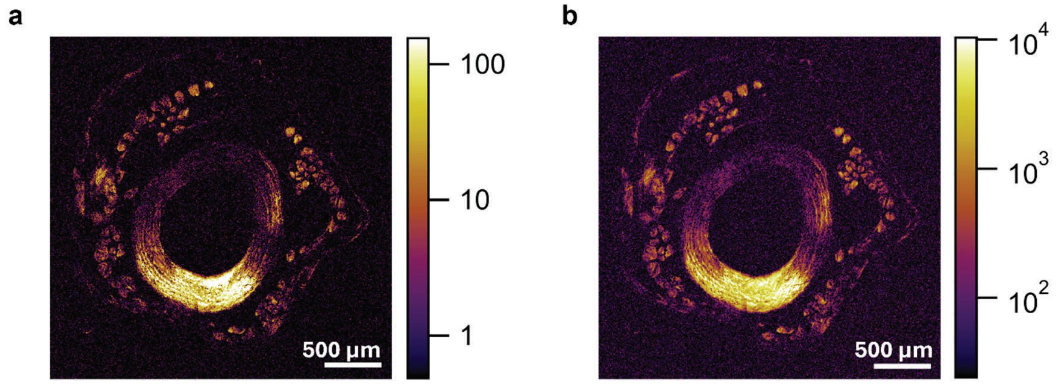


**Supplementary Fig. S3 | Millimeter-scale imaging of mouse tail annulus fibrosus**

a,b, Integrated coherent- and incoherent-component images, respectively, reconstructed from a single hyperspectral acquisition over a 3 × 3 mm field in mouse tail annulus fibrosus. Intensities are displayed on logarithmic color scales. Scale bars, 500 μm. Both components were recovered throughout the scan, and the images delineated the collagen fascicle architecture. Because the sampling pitch was coarser than in the muscle and dermis measurements, this dataset was excluded from quantitative spatial-frequency comparisons and was used only to demonstrate large-area spectral channel separation.

**Supplementary Note 4 | Effect of Transverse-Wavevector Correlations on Walk-Off-Induced Effective Phase Mismatch**

We discuss the possibility that the transverse-wavevector correlations of twin beams can reduce the effective phase mismatch in the SFG process arising from walk-off, thereby increasing the effective interaction length.

Here, we consider an SFG process in a uniform birefringent medium. In biological samples such as collagen, which are the focus of this study, the refractive-index distribution and fiber orientation vary spatially and are considerably more complex. The essential point of interest here is whether the transverse-wavevector correlations of twin beams can reduce the effect of the relative phase shift arising from walk-off associated with refractive-index anisotropy. This relative phase shift gives rise to an effective phase mismatch in the SFG process. By considering a simplified model based on a uniform birefringent medium, we expect to clarify the role of the transverse-wavevector correlations of twin beams in this process.

Specifically, we consider a type-I SFG process under birefringent phase matching, in which two ordinarily polarized input fields are up-converted into an extraordinarily polarized SFG field. This configuration involves spatial walk-off of the extraordinary SFG field. Theoretical analyses of SFG with twin beams have predicted that the influence of spatial walk-off can be reduced for the coherent SFG component generated from correlated mode pairs [1]. In addition, SFG experiments using phase-conjugate twin beams have demonstrated a coherent SFG component with high contrast against the incoherent background, in agreement with this theoretical picture [2]. Furthermore, experiments using the twin-beam source employed in the present study have also demonstrated high-contrast coherent upconversion, suggesting that a similar mechanism may contribute to the observed behavior [3].

According to Ref. [4], the longitudinal wavevector components can be expanded as

$$k_{jz}(q_j, \delta\omega_j) \simeq k_j + k_j' \delta\omega_j + \frac{1}{2} k_j'' \delta\omega_j^2 - \frac{|\, q_j \,|^2}{2k_j},\ j = s, i,$$

for the ordinary input fields, and

$$k_{\mathrm{SFG},z}(\boldsymbol{q}_{\mathrm{SFG}}, \delta\omega_{\mathrm{SFG}}) \simeq k_{\mathrm{SFG}} + k_{\mathrm{SFG}}' \delta\omega_{\mathrm{SFG}} + \frac{1}{2} k_{\mathrm{SFG}}'' \delta\omega_{\mathrm{SFG}}^2 + \frac{\partial k_{\mathrm{SFG}}}{\partial q_{\mathrm{SFG},x}} q_{\mathrm{SFG},x} - \frac{|\, q_{\mathrm{SFG}} \,|^2}{2k_{\mathrm{SFG}}},$$

for the extraordinary SFG field. Here, the notation in Ref. [4] has been rewritten using the notation adopted in the present work. $\delta\omega_s$, $\delta\omega_i$, and $\delta\omega_{\mathrm{SFG}}$ denote the frequency detunings from the respective central frequencies. The $x$ axis is taken along the walk-off direction, and $\frac{\partial k_{\mathrm{SFG}}}{\partial q_{\mathrm{SFG},x}}$ corresponds to the walk-off angle.

A key difference between the longitudinal wavevector components of the ordinary input fields and the extraordinary SFG field is that the latter contains the term $\frac{\partial k_{\mathrm{SFG}}}{\partial q_{\mathrm{SFG},x}} q_{\mathrm{SFG},x}$. This difference gives rise to a transverse-wavevector-dependent relative phase shift between the ordinary input fields and the extraordinary SFG field during propagation. The resulting phase shift can therefore be regarded as a walk-off-dependent contribution to the effective phase mismatch of the SFG process.

For correlated signal–idler mode pairs in twin beams, the transverse wavevectors ideally satisfy $\boldsymbol{q}_s + \boldsymbol{q}_i = 0$. By transverse momentum conservation, this gives $\boldsymbol{q}_{\mathrm{SFG}} = 0$, for the coherent SFG component, and therefore the walk-off-induced term $\frac{\partial k_{\mathrm{SFG}}}{\partial q_{\mathrm{SFG},x}} q_{\mathrm{SFG},x}$ vanishes. Thus, the walk-off-induced phase contribution can be eliminated for the coherent SFG component, thereby increasing its effective interaction length.
